# Distinct Membrane Binding Properties of the Two Non-visual Arrestins

**DOI:** 10.1101/2025.03.06.641943

**Authors:** Thomas D. Killeen, Katelyn Tepper, Kyle W. Miller, Yasmin Aydin, Ya Zhuo, Shuaitong Zhao, Jason M. Conley, Rachel Tat, Candice Klug, Adriano Marchese, Valerică Raicu, Qiuyan Chen

## Abstract

Membrane interactions play a crucial role in regulating arrestin activation and its binding to phosphorylated G protein-coupled receptors (GPCRs). Here, we utilize *in vitro* biophysical approaches and cell-based fluorescence intensity fluctuation analysis to systematically compare the membrane-binding properties of the two highly conserved arrestin subtypes, arrestin-2 and arrestin-3, under basal and stimulated conditions. Our findings reveal that arrestin-2 selectively engages the PI(4,5)P_2_-containing nanodiscs via its C-edge, whereas arrestin-3 primarily utilizes its finger loop to interact with negatively charged lipids. Notably, while the lipid bilayer alone does not activate arrestin, it synergistically enhances arrestin-2/3 activation in conjunction with a phosphorylated GPCR C-tail. Additionally, the spacing between receptor phosphorylation sites and the lipid bilayer modulates arrestin−membrane assembly. Live cell tracking further demonstrates that arrestin-2 and arrestin-3 exhibit distinct plasma membrane dissociation dynamics. These findings provide novel insights into the mechanisms governing arrestin activation and its functional interplay with membranes.

## Introduction

Two ubiquitously expressed arrestins, arrestin-2 (Arr2) and arrestin-3 (Arr3) (also known as β-arrestin1 and β-arrestin2, respectively), regulate signaling of hundreds of G protein-coupled receptors (GPCRs)^1,2^. Decades of research indicate that arrestin activation and its downstream signaling roles are primarily determined by interactions with the receptor’s phosphorylated C-tail^3–5^, intracellular loops^6,7^, and transmembrane (TM) core^8–10^. However, emerging evidence suggests that membrane association is also an important factor in modulating arrestin signaling, adding a new dimension to our understanding of these multifunctional proteins^11–17^. For instance, Janetzko *et al.* demonstrated that membrane phosphoinositides regulate the assembly and dynamics of GPCR−arrestin complexes^13^. A single-molecule microscopy study by Grimes *et al.* revealed that arrestin associates with the plasma membrane both before and after receptor stimulation, and membrane binding facilitates its interactions with receptors^11^.

Structural evidence further underscores the importance of membrane interactions in arrestin function. Both Arr2 and Arr3 harbor a phosphoinositide-binding site on the concave side of their C-domain^16,18,19^. All cryo-EM structures of GPCR−Arr2 complexes determined to date show that the C-edge of Arr2 directly inserts into the detergent micelles or nanodiscs surrounding GPCRs, contributing to stabilization of the complex^16,20–24^. Our recent findings demonstrate that both Arr2 and Arr3 utilize their finger loop—previously identified as the "activation sensor" for its role in engaging the receptor TM core—to interact with detergent micelles when in complex with the atypical chemokine receptor 3 (ACKR3)^24^. Notably, the C-edge of Arr3 lacks a defined membrane-binding segment, leading to increased dynamics of the C-domain when complexed with ACKR3 compared to Arr2^24^. This observation suggests that the membrane binding properties of Arr2 and Arr3 differ. Despite these advances, it remains unclear how the structural elements in arrestins coordinate to bind lipid bilayers in the basal and activated states.

Different phosphorylation patterns, or "barcodes" installed by distinct GPCR kinase (GRK) subtypes, have been reported to lead to unique functional outcomes^3,4,25^. Our recent finding that the spacing between phosphorylation sites and the TM core of ACKR3 governs differences in its assembly with Arr2 and Arr3^24^ broadens the potential mechanisms by which phosphorylation barcodes can regulate arrestin signaling. It remains unclear whether the regulatory effects of spacing are independent of the ACKR3 receptor core and whether this phenomenon extends to other GPCRs.

To this end we systematically compared the membrane-binding properties of Arr2 and Arr3 in various functional states. Our results demonstrate that Arr3, but not Arr2, primarily relies on its finger loop for lipid binding in the basal state, and that activation enhances Arr2 binding to PI(4,5)P_2_ but reduces Arr3 binding to this lipid. Using lipidated phosphopeptides derived from the ACKR3, we further demonstrate that membrane association and phosphopeptide binding synergistically activate Arr2 and Arr3 and that the spacing between phosphorylation sites and the lipid tether modulates arrestin−membrane assembly. Finally, live-cell fluorescence intensity fluctuation (FIF) analysis confirmed that the dynamics of Arr2 and Arr3 in membrane association and dissociation vary in native membranes and clathrin-coated pits (CCPs) upon activation of the M_2_ muscarinic acetylcholine receptor (M_2_R).

### Arr2 and Arr3 Bind to Membrane via Different Structural Elements

To mimic the lipid bilayer, we prepared four types of nanodiscs with distinct lipid compositions: 1) 100% 1-palmitoyl-2-oleoyl-glycero-3-phosphocholine (POPC); 2) 40% 1-palmitoyl-2-oleoyl-sn-glycero-3-phospho-L-serine (POPS) with 60% POPC (molar ratio); 3) 40% 1-palmitoyl-2-oleoyl-sn-glycero-3-phospho-(1’-rac-glycerol) (POPG) with 60% POPC; and 4) 10% L-α-phosphatidylinositol-4,5-bisphosphate (PI(4,5)P_2_) with 90% POPC (**Fig. 1A**). The scaffolding protein we used forms nanodiscs with a diameter of approximately 9 nm (NW9)^26^ and includes a His tag for pulldown assays.

**Figure 1.**
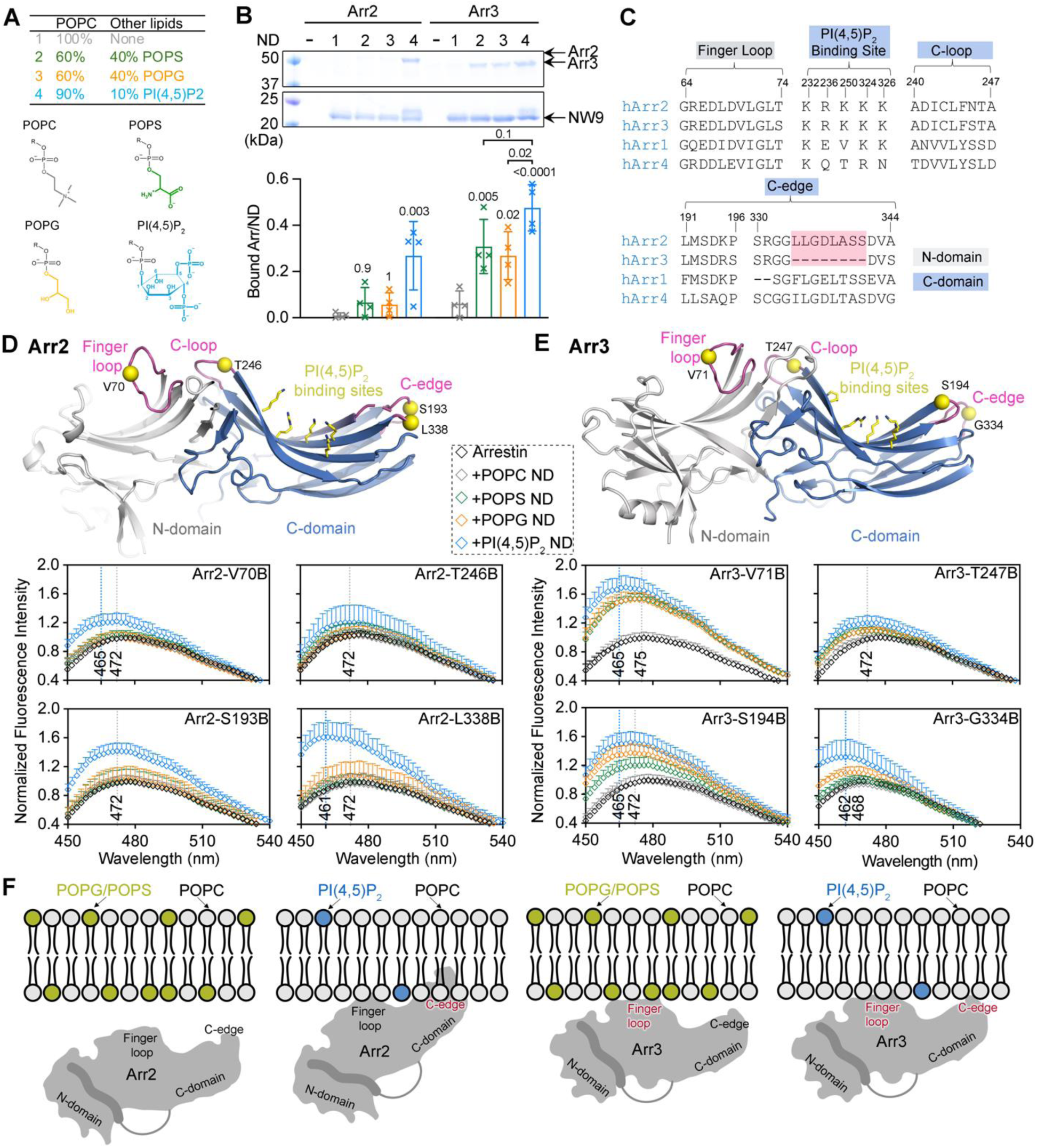
Distinct membrane binding modes of basal Arr2 and Arr3. **A.** The lipid composition of four nanodiscs used for this study. The headgroup structures of the four lipids are shown. **B.** A His-pulldown assay shows that Arr2 selectively binds to the PI(4,5)P_2_ nanodiscs, whereas Arr3 is less selective and binds to the POPG, POPS, or PI(4,5)P_2_ nanodiscs. One-way ANOVA followed by a Tukey multiple comparison test was used to compare bound arrestin to nanodisc ratios within the Arr2 and Arr3 group, respectively. Error bars represent S.D. from three technical replicates. The *p* value comparing its binding to the POPC nanodiscs is shown on top of the column. **C.** Sequence alignments of potential membrane-binding sites on arrestins. The 338 C-edge loop, which Arr3 lacks, is shown in pink. **D, E**. The crystal structure of Arr2 in the basal state (PDB entry 1G4M) (**D**) and Arr3 in the basal state (PDB entry 3P2D) (**E**) with the Cα of the four labelled sites shown as yellow spheres and the PI(4,5)P_2_ binding sites shown as yellow sticks. The fluorescence spectra of Arr2 (**D**) and Arr3 (**E**) variant in the absence (black) or presence of the POPC (grey), POPS (green), POPG (orange) or PI(4,5)P_2_ (blue) nanodiscs. (**F**) Cartoon mode showing distinct modes of Arr2 and Arr3 binding to the lipid bilayer containing POPG/POPS or PI(4,5)P_2_.

We first assessed how Arr2/3 proteins bind to these four types of nanodiscs in their basal state. The pulldown results revealed that Arr2 selectively binds to the PI(4,5)P_2_ nanodiscs, with negligible binding to the POPC, POPS, or POPG nanodiscs (**Fig. 1B**). In contrast, Arr3 interacts with the PI(4,5)P_2_ nanodiscs, as well as with the POPS and POPG nanodiscs, while its binding to the POPC nanodiscs remains minimal (**Fig. 1B**). Notably, Arr3 binding to the PI(4,5)P_2_ nanodiscs is approximately 50-60% higher than its binding to the POPS or POPG nanodiscs, suggesting that the PI(4,5)P_2_ headgroups are more efficient at promoting Arr3 interaction with lipid bilayer.

To understand the distinct membrane-binding properties of these two highly homologous arrestin subtypes, we next investigated the structural elements on Arr2/3 involved in lipid binding. Previous studies show that the C-domain of Arr2/3 contains several membrane-anchoring elements: the PI(4,5)P_2_ binding sites^16,27^, the C-loop^11^ and the C-edge^11,16,22,23^(**Fig. 1C**). Our recent study reveals that the N-domain of Arr2/3 also contributes to membrane binding via the finger loop^24^(**Fig. 1C**). Those sites are conserved, except that Arr3 lacks an extended membrane-anchoring segment (residues 334-341, **Fig. 1C**). To investigate the functional roles of these elements, we attached a bimane fluorophore to four positions on Arr2 (**Fig. 1D**): the finger loop (V70), the C-loop (T246), and the C-edge (S193 and L338). Similarly, we labeled the analogous positions on Arr3 (**Fig. 1E**): the finger loop (V71), the C-loop (T247), and the C-edge (S194 and G334). The bimane fluorophore, which is sensitive to its chemical environment, was used to monitor arrestin interactions with lipids^12,13^, phosphopeptides^13^, and GPCRs^12,22,24,28^.

Consistent with the pulldown results (**Fig. 1B**), we detected substantial fluorescence changes for Arr2 only in the presence of the PI(4,5)P_2_ nanodiscs (**Fig. 1D**), confirming its high selectivity for PI(4,5)P_2_. The largest fluorescence changes were observed at the S193B and L338B sites, consistent with the C-edge moving into a more hydrophobic environment (**Fig. 1F**). Bimane fluorescence at the finger loop (V70B) exhibited a slight increase in intensity (1.2±0.1-fold) and a peak shift (from 472 to 465 nm), supporting its role in membrane binding. In contrast, bimane fluorescence at the C-loop did not show any significant changes. Collectively, these results suggest that PI(4,5)P_2_ recruits the C-domain of Arr2 to the lipid bilayer, promoting the C-edge binding, and to a lesser extent, enabling the finger loop anchoring to the membrane (**Fig. 1F**).

For Arr3, the finger loop (V71C), rather than the C-edge, showed the most pronounced response to the PI(4,5)P_2_ nanodiscs, with both an increase in intensity (1.7±0.2-fold) and a peak shift (from 475 to 465 nm) (**Fig. 1E**). Similar fluorescence changes in the finger loop were also observed in the presence of the POPG and POPS nanodiscs. This indicates that in the basal state, the finger loop of Arr3 anchors to the membrane in the presence of negatively charged lipids (**Fig. 1F**). In contrast, the POPS and POPG nanodiscs induced less or no fluorescence changes at the S194B and G334B sites compared to the PI(4,5)P_2_ nanodiscs (**Fig. 1E**). This suggests that PI(4,5)P_2_ binding brings the C-domain, including the C-edge, closer to the lipid bilayer. However, unlike Arr2, the Arr3 334 C-edge cannot stably anchor to the membrane, even when in close proximity, due to the absence of a membrane-binding loop (**Fig. 1C**).

Collectively, these results show that Arr2 selectively binds to the PI(4,5)P_2_ nanodiscs primarily via its C-edge, whereas Arr3 anchors to negatively charged lipids mainly via its finger loop in the N-domain. Additionally, PI(4,5)P_2_ enhances the interaction between the Arr3 C-domain and the membrane (**Fig. 1F**).

### Contrasting PI(4,5)P_2_ Binding Profiles of Pre-activated Arr2 and Arr3 Compared to Their Respective WT Forms

The activation of arrestins involves a rotation between the N- and C-domains, which can realign the various membrane-binding elements and alter their interaction with lipids. We therefore examined how the pre-activated Arr2/3 (Arr2*: I386A, V387A, F388A; Arr3*: I387A, V388A, F389A) interact with nanodiscs containing different lipids. These triple alanine mutations destabilize the hydrophobic interactions anchoring the arrestin C-terminus to the N-domain, thereby releasing this autoinhibitory intramolecular interaction and allowing arrestins to relax into their activated states^29^.

We first compared the binding properties of Arr2* and Arr3* to their respective wild-type (WT) forms. Pulldown results showed that Arr2* selectively binds to the PI(4,5)P_2_ nanodiscs, similar to WT Arr2. The amount of bound Arr2* increased by ∼60% compared to WT Arr2 (**Fig. 2A**). In contrast, Arr3* binding to the PI(4,5)P_2_ nanodiscs was reduced by ∼50% compared to WT Arr3, reaching a level comparable to its binding to the POPS and POPG nanodiscs (**Fig. 2B)**. This result is unexpected, as one key role of arrestins is to direct activated GPCRs to CCPs where PI(4,5)P_2_ is enriched during the early formation stage^30^. Therefore, we anticipated that both arrestins would exhibit enhanced binding to PI(4,5)P_2_ upon activation.

**Figure 2.**
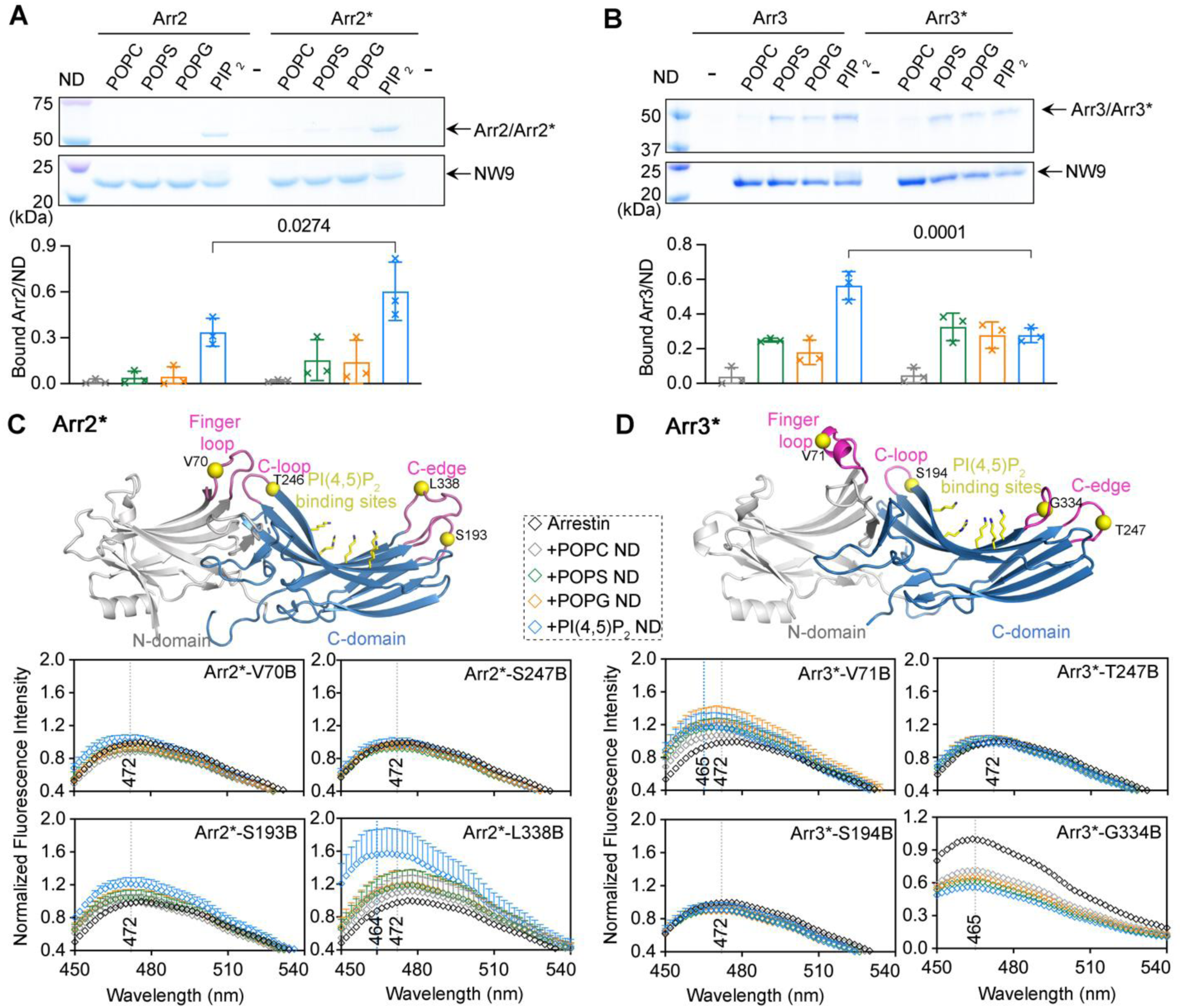
Distinct membrane binding modes of Arr2* and Arr3*. **A.** A His pulldown assay shows that Arr2* selectively binds to the PI(4,5)P_2_ nanodiscs and the binding is ∼60% higher than that of WT Arr2. **B**. A His pulldown assay shows that Arr3* binds to the POPS-, POPG- and PI(4,5)P_2_ nanodiscs but the binding to the PI(4,5)P_2_ nanodiscs is ∼50% lower than that of WT Arr3. **C, D.** The crystal structure of active Arr2 (PDB entry 4JQI) (**C**) and active Arr3 (PDB entry 5TV1) (**D**) with the four labeled sites shown as yellow spheres and the PI(4,5)P_2_ binding sites shown as yellow sticks. The fluorescence spectra of Arr2* variants (**C**) or Arr3* variants (**D**) in the absence (black) or presence of the POPC (grey), POPS (green), POPG (orange) or PI(4,5)P_2_ (blue) nanodiscs.

To further investigate the contrasting PI(4,5)P_2_ binding profiles of Arr2* and Arr3*, we labeled the same positions in Arr2* and Arr3* as in their WT counterparts with the bimane fluorophore (**Fig. 2C, D vs. Fig. 1D, E**). In Arr2*, only the 338 C-edge bimane showed a substantial fluorescence change in the presence of the PI(4,5)P_2_ nanodiscs, with the magnitude of the change comparable to that of Arr2 (**Fig. 2C**). We then titrated the PI(4,5)P_2_ nanodiscs against Arr2-L338B and Arr2*-L338B and estimated the binding affinity (K_d_)(**Extended Data Fig. 1A**). The affinity of Arr2* (K_d_=20±20 nM) was significantly higher than that of Arr2 (K_d_=300±40 nM), consistent with the increased binding of Arr2* to the PI(4,5)P_2_ nanodiscs observed in pulldowns (**Fig. 2A**). Interestingly, neither the 193 C-edge loop nor the finger loop in Arr2* showed fluorescence changes (**Fig. 2C**), suggesting Arr2* primarily anchors to the PI(4,5)P_2_ nanodiscs via the 338 C-edge loop and the nearby PI(4,5)P_2_ binding site.

In case of Arr3*, the finger loop (V71B) exhibited much smaller changes compared to WT Arr3, and the C-edge (G334B) showed decreased fluorescence changes (**Fig. 2D**), in contrast to the increased changes observed in basal Arr3 (**Fig. 1E**). This indicates that Arr3* adopts a conformation distinct from Arr3 when engaging the membrane, likely leading to its reduced binding to the PI(4,5)P_2_ nanodiscs, as observed in the pulldown assay (**Fig. 2B**). This could facilitate Arr3* dissociation from the membrane after detaching from the activated GPCR in the cell.

### Membrane Association Does Not Induce Arr2 Activation

Recent studies reported that a soluble PI(4,5)P_2_ derivative, diC8-PI(4,5)P_2_, promotes conformational changes consistent with Arr2 and Arr3 activation but does not induce the full release of their autoinhibitory C-terminus^13,31^. Since Arr2 contains multiple structural elements that interact with the lipid bilayer beyond PI(4,5)P_2_ (**Fig. 1D**), we examined how different nanodiscs influence the C-terminal release and consequently, the activation of Arr2.

We introduced a pair of cysteines in Arr2 (A12C/A392C), where A12C is located on the β1-strand of the N-domain and A392C is at the C-terminus of Arr2 (**Fig. 3A**). These cysteines were labeled with spin labels, and the double electron-electron resonance (DEER) was used to directly measure the distance between these two positions (**Fig. 3A**). In the basal state, the primary distance is centered around 18 Å, as the C-terminus of arrestin is attached to its N-domain. Activation by p-Rho* resulted in a longer distance of approximately 62 Å, consistent with the release of the C-terminus (**Fig. 3B**). All nanodiscs induce very subtle changes in DEER distance distribution, with the main population remaining in the basal state, centered around 18 Å (**Fig. 3C**). Although Arr2 exhibits highly selective binding to the PI(4,5)P_2_ nanodiscs (**Fig. 1A**), the PI(4,5)P_2_ nanodiscs does not promote its activation, similarly to all other types of nanodiscs (**Fig. 3C**).

**Figure 3.**
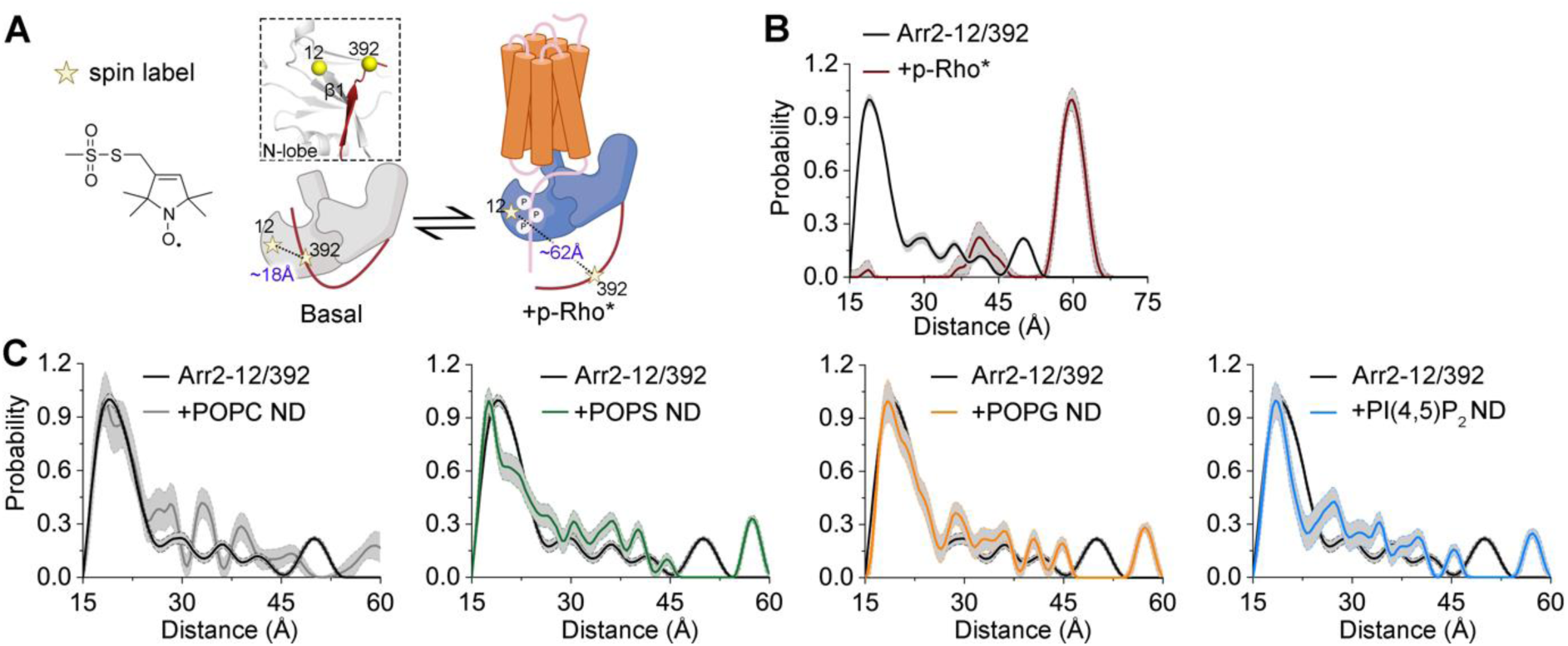
Membrane association does not trigger the activation of Arr2. **A.** Schematic illustrating how the DEER distance change corresponds to the relative position between the Arr2 C-terminus and N-domain. The inset shows the positions of the A12C and A392C pair on Arr2 (PDB entry 1G4M), with the Cα atoms shown as yellow spheres. **B, C.** DEER distance distribution of Arr2-12/392 in the absence or the presence of p-Rho* (**B**) or the POPC, POPS, POPG, and PI(4,5)P_2_ nanodiscs (ND) (**C**).

### Phosphopeptides and Lipid Bilayer Synergistically Activate Arrestins

Our recent study shows that both Arr2 and Arr3 engage ACKR3 primarily by interacting with its phosphorylated C-tail and anchoring to the surrounding detergents, without extensive contact with the cytoplasmic cleft of the receptor^24^. This finding prompted us to investigate whether the membrane and phosphorylated receptor C-tail are sufficient to activate Arr2 and Arr3.

To test this, we synthesized four peptides based on the unique GRK5 phosphorylation signature on ACKR3^32^ (**Fig. 4A**) with N-terminal palmitoyl to tether them to the lipid bilayer (**Fig. 4B**). ACRK3_pp1_ and ACRK3_pp2_ feature both palmitoylation and phosphorylation, with linkers of 5 (ACKR3_pp1_) or 17 (ACKR3_pp2_) residues between the palmitoylation site and the first phospho-Ser residue, respectively. These designs mimic the GRK5 and GRK2 phospho-barcodes of ACKR3^32^ and allow us to assess how linker length affects arrestin activation and its interaction with lipids. ACRK3_pp3_ contains all phosphorylation sites but lacks the palmitoylation, whereas ACKR3_p4_ is palmitoylated but has no phosphorylation.

**Figure 4.**
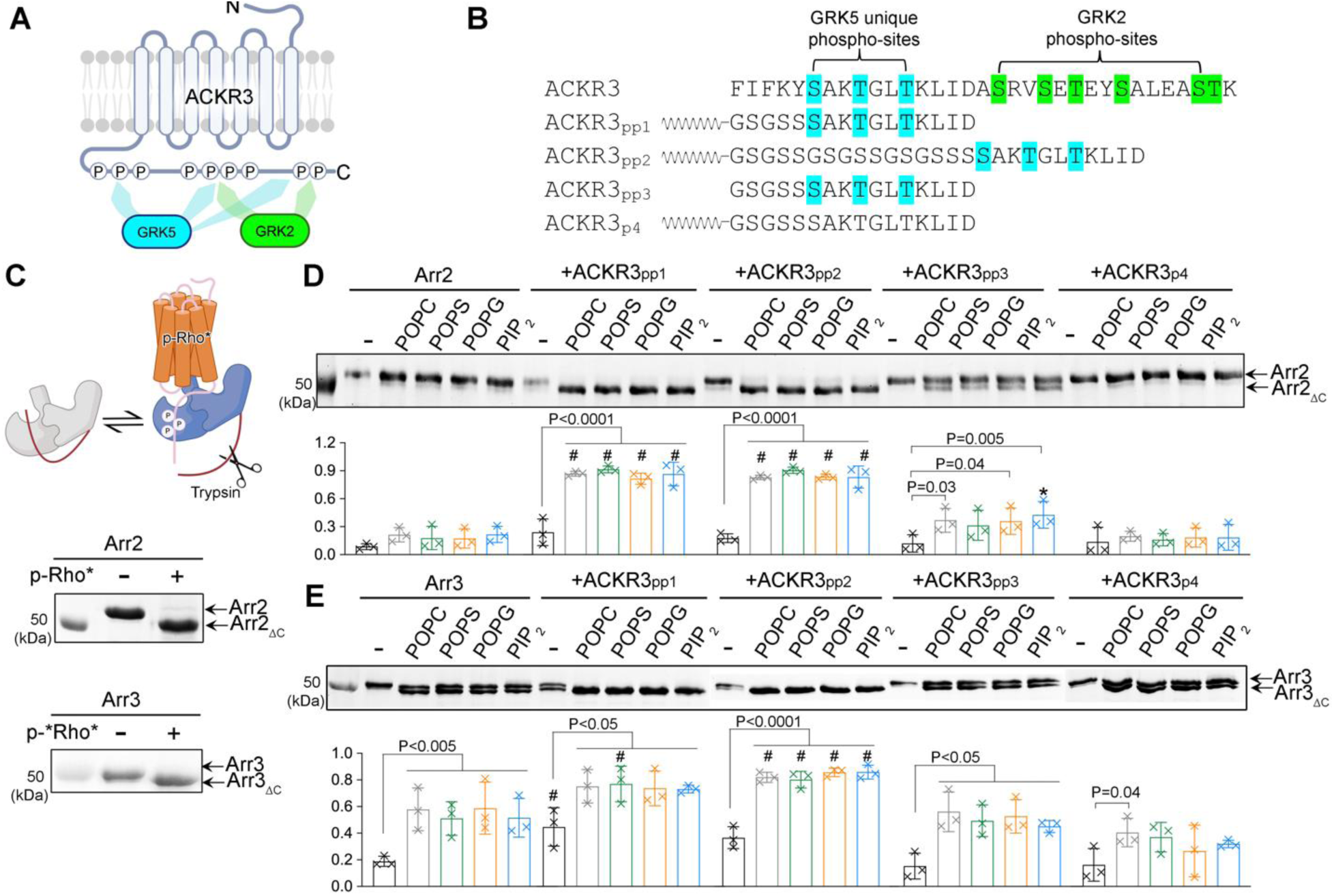
Phosphopeptides and lipids synergistically activate Arr2 and Arr3. **A.** Diagram of ACKR3 highlighting GRK5 and GRK2 phosphorylation sites. GRK5 phosphorylates both the proximal and distal sites on ACKR3, whereas GRK2 only targets distal sites. **B.** Table listing the sequences of ACKR3 C-tail and the four peptides used in this study. Unique GRK5 phosphorylation sites are shown in blue and GRK2 phosphorylation sites are shown in green. **C.** Schematic illustrating how the trypsin digestion pattern correlates with the release of arrestin C-terminus upon p-Rho* binding. The gel shows that a truncated form of Arr2 (Arr2_ΔC_) and Arr3 (Arr3_ΔC_) is produced in the presence of p-Rho* indicating the activation of Arr2 and Arr3, respectively. **D, E.** The trypsin digestion patterns of Arr2 (**D**) or Arr3 (**E**) in the absence and presence of four nanodisc types and four ACKR3-derived peptides listed in B.

We first examined the effects of nanodiscs and peptides on arrestin activation using a limited trypsin digestion assay (**Fig. 4C**). P-Rho* served as a positive control, as it induces the release of Arr2 or Arr3 C-terminus^33^, exposing additional lysines for trypsin digestion that generates truncated forms of Arr2 (Arr2_ΔC_) and Arr3 (Arr3_ΔC_) (**Fig. 4C**).

Neither nanodiscs nor peptides alone activated Arr2 (**Fig. 4D**). However, nanodiscs with either ACKR3_pp1_ or ACKR3_pp2_ induced nearly 100% activation. Both the lipid tether and the receptor C-tail phosphorylation were essential for the full activation of Arr2, as ACRK3_pp3_ activated only ∼30% of Arr2 and ACKR3_p4_ failed to trigger any activation. Notably, the phospholipid composition of the nanodiscs did not affect activation, indicating that Arr2 requires a lipid bilayer rather than specific lipid headgroups for activation, provided a phosphorylated receptor C-tail is nearby. Similar results were observed for Arr3 (**Fig. 4E**), although nanodiscs or peptides alone partially activated Arr3, suggesting that Arr3 is more readily activated than Arr2.

Collectively, our results demonstrate that phosphorylated receptor C-tails and lipid bilayer synergistically activate Arr2 and Arr3, emphasizing the critical interplay between membrane anchoring and phosphopeptide binding in arrestin activation.

### Membrane Tethered Phosphopeptides Modulate Arrestin−Membrane Assembly

We next tested how different peptides influence arrestin interaction with the nanodiscs. Pull-down assays showed that both ACKR3_pp1_ and ACKR3_pp2_ enhanced Arr2/3 binding to the POPS and POPG nanodiscs, reaching levels comparable to those observed with the PI(4,5)P_2_ nanodiscs. Additionally, ACKR3_pp1_ increased Arr2/3 binding to the POPC (**Extended Data Fig. 1B**). In contrast, neither ACKR3_pp3_ nor ACKR3_p4_ affected Arr2 or Arr3 binding. These findings suggest that membrane-tethered phosphopeptides promote Arr2/3 engagement with lipid bilayers without a specific requirement for PI(4,5)P_2_. This is consistent with previous observations that the phosphorylated C-tail of GPCRs alone is sufficient to retain arrestins at the membrane^9,34–36^.

Our recent study shows that the spacing between phosphorylation sites and the TM core of ACKR3 dictates how the receptor complex assembles with Arr2 or Arr3^24^. On the ACKR3 C-tail, GRK5 phosphorylation sites are located 14 residues closer to the TM core than GRK2 phosphorylation sites (**Fig. 4A**). This proximity constrains the interaction with arrestins, resulting in the formation of more rigid complexes with a more extensive interface involving the intracellular surface of the TM core and the surrounding detergent^24^. This raises the intriguing question of whether ACKR3_pp1_ and ACKR3_pp2_, which mimic the GRK5 and GRK2 phosphorylation barcodes via their differing spacer lengths, exhibit similar effects on arrestin-membrane interaction.

To test this, we examined the structural elements of Arr2 and Arr3 involved in binding in the presence of different peptides, using our bimane-labeled variants. Lower concentrations of Arr2/3 and nanodiscs were used in this assay (0.5 μM in Fig. 5 vs. 2 μM in Fig. 1D) to reduce the signal change caused solely by lipid binding (**Fig. 5A**). ACKR3_pp1_ and ACKR3_pp2_ induced comparable fluorescence changes in the Arr2 finger loop (V70) in the presence of the POPG or PI(4,5)P_2_ nanodiscs (**Fig. 5A**). Both the lipid tether and phosphorylation are essential for finger loop engagement, because neither ACKR3_pp3_ nor ACKR3_p4_ caused fluorescence changes at the V70 position (**Fig. 5A**). Interestingly, ACKR3_pp1_ induced greater fluorescence changes in both C-edge positions compared to ACKR3_pp2_ (**Fig. 5A**), consistent with a shorter linker holding Arr2 closer to the lipid bilayer, forcing the C-edge to anchor more strongly (**Fig. 5B**). This observation aligns with our previous finding that Arr2 makes more extensive contacts with ACKR3 and solubilizing micelles when its proximal sites are phosphorylated by GRK5 compared to when its distal sites are phosphorylated by GRK2^24^. Additionally, PI(4,5)P_2_ likely provides an auxiliary anchoring site on the Arr2 C-domain, further stabilizing its interaction with the membrane (**Fig. 5A**; **blue vs. orange curve**).

**Figure 5.**
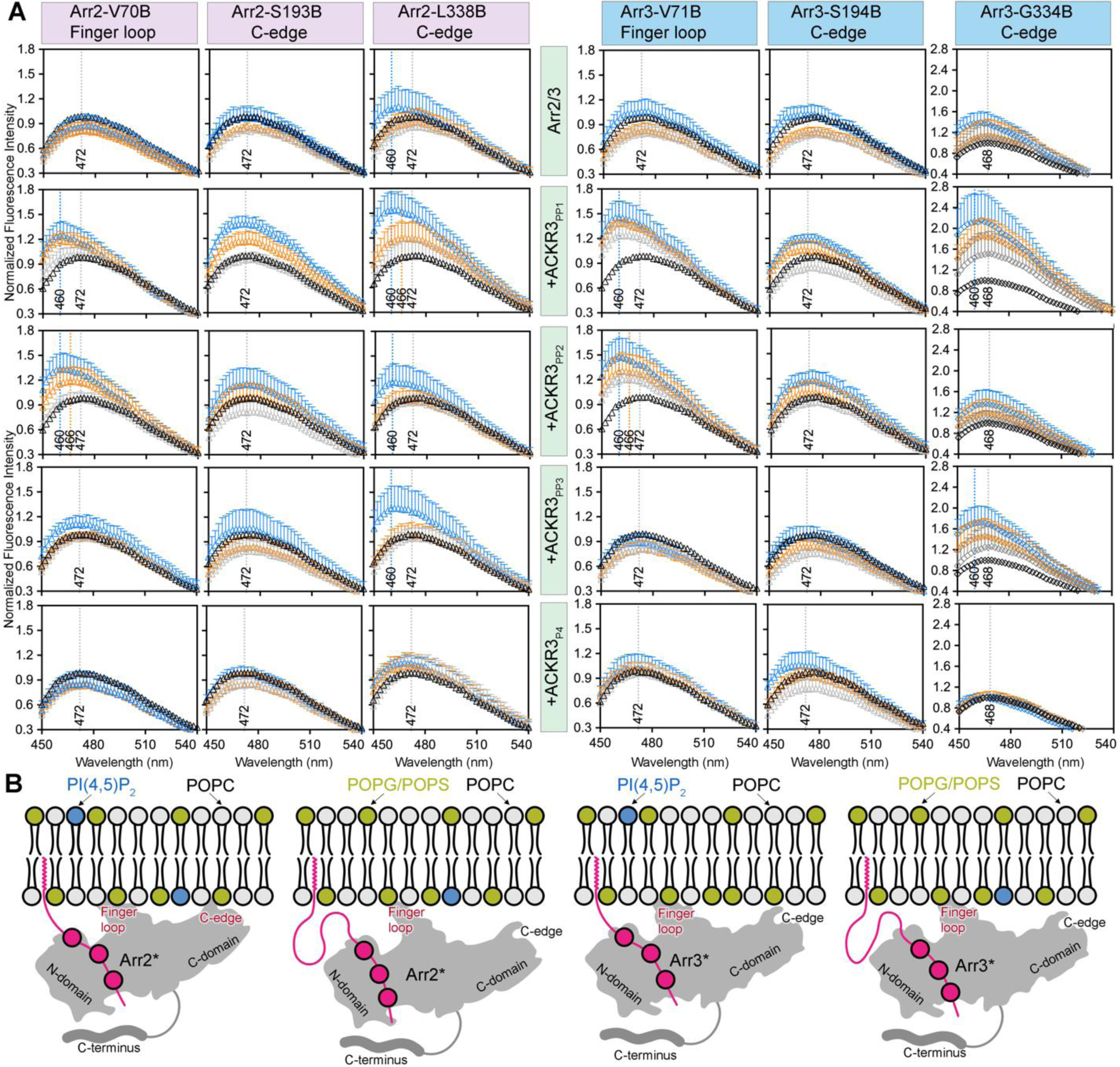
Membrane Association of Arr2/3 is Regulated by Membrane-Tethered Phosphopeptides. **A.** Fluorescence spectra of Arr2 and Arr3 in the absence (black) or presence of the POPC (grey), POPG (orange) or PI(4,5)P_2_ (blue) nanodiscs, with and without ACKR3_pp1_, ACKR3_pp2_, ACKR3_pp3_ Or ACKR3_p4_. The fluorescence spectra in the presence of the POPS nanodiscs were similar to the POPG nanodiscs and are not shown. **B.** Cartoon modes illustrating distinct modes of Arr2 and Arr3 binding to the membrane in the presence of ACKR3_pp1_ and ACKR_pp2_.

Arr3 exhibited similar responses to phosphopeptides and nanodiscs as Arr2. Both ACKR3_pp1_ and ACKR3_pp2_, but not ACKR3_pp3_ or ACKR3_pp4_, induced substantial changes in the Arr3 finger loop upon binding to all nanodisc types. Notably, these changes are more pronounced in Arr3 compared to Arr2 (**Fig. 5A**). ACKR3_pp1_ triggered greater fluorescence changes in the Arr3 334 C-edge than ACKR3_pp2_ (**Fig. 5A**), suggesting that a shorter linker also brings Arr3 C-domain closer to the membrane. However, because the Arr3 334 C-edge lacks a membrane-anchoring segment (**Fig. 1C**), it is unable to stably anchor to the lipid bilayer even when in close proximity. Our previous findings suggest that the Arr3 C-domain exhibits higher dynamics in complex with ACKR3 than that of Arr2^24^.

Taken together, these findings suggest that the phosphorylated receptor C-tail pulls the Arr2 and Arr3 N-domain close to the membrane, promoting the finger loop engagement with either the membrane or the receptor core, thereby pivoting the N-domain towards the lipid bilayer. C-edge involvement in membrane binding depends on the spacer length between phosphorylation sites and the membrane but is limited in Arr3 due to the absence of a membrane-anchoring segment.

### Arr2 and Arr3 Exhibit Distinct Membrane Interaction Dynamics in Cells

We next compared how Arr2 and Arr3 translocate to the plasma membrane and CCPs upon receptor activation in cells. The M_2_ muscarinic acetylcholine receptor (M_2_R) was selected due to its strong colocalization with arrestins in PI(4,5)P_2_-enriched CCPs upon activation^37,38^. To track M_2_R and Arr2/3, fluorescent proteins mEGFP and mCitrine were fused to the N-terminus of M_2_R and Arr2/3, respectively. HEK293 cells were transiently transfected with both constructs and imaged 24-48 hours post-transfection using a two-photon fluorescence micro-spectroscope^39^. Imaging scans were performed at the basolateral membrane before and 2, 10, 20, 30, 40, and 50 minutes after stimulation with the M_2_R agonist, carbachol (**Fig. 6A**). Using fluorescence intensity fluctuation (FIF) analysis (**Extended Data Fig. 2**)^40^, we determined the Arr2/3-to-M_2_R ratio at each time point at the plasma membrane (**Fig. 6B**) and within the CCPs (**Fig.6C**).

**Figure 6.**
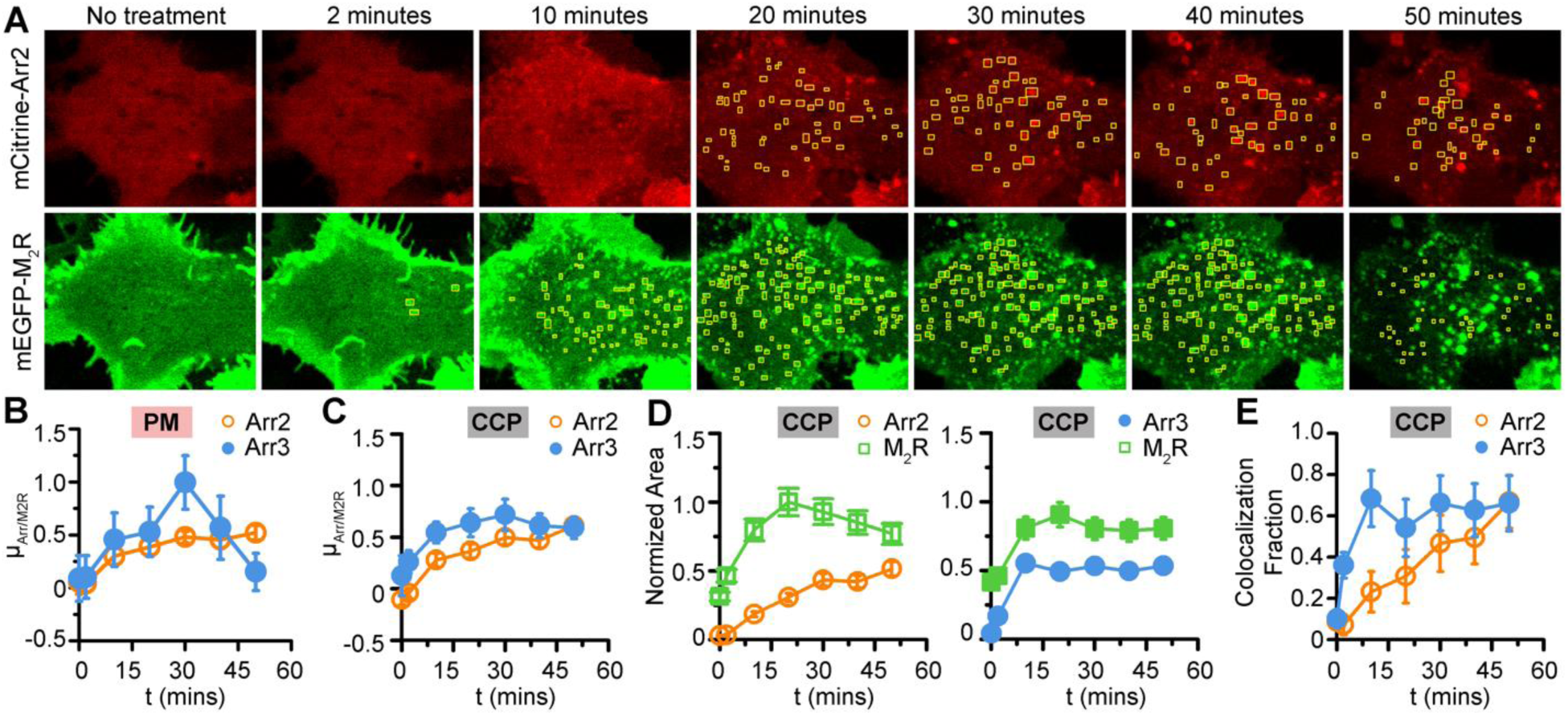
Arr2 and Arr3 Exhibit Distinct Interaction Dynamics with Plasma Membrane and CCPs. **A**. Zoomed-in view of a single cell expressing both M_2_R and Arr2 before and after carbachol stimulation. CCPs were visually identified as distinct circular structures with diameters of several hundred nanometers and highlighted with yellow rectangle boxes. **B, C.** Distributions of Arr2-to-M_2_R and Arr3-to-M_2_R concentration ratios at selected time points at the plasma membrane (PM) (**B**) and within the CCPs (**C**). **D.** Area of CCPs containing Arr2/3 or M_2_R plotted against treatment time. **E.** Fraction of CCPs containing both M_2_R and Arr2/3 as a proxy for colocalization estimation, plotted against treatment time.

No pre-association with the plasma membrane was observed for either Arr2 or Arr3 (**Fig. 6B**). During the first 30 minutes post-stimulation, both Arr2 and Arr3 showed a gradual increase in membrane association, with Arr3 reaching a higher peak than Arr2 (**Fig. 6B**). Notably, after 30 minutes, the Arr2-to-M_2_R ratio stabilized, whereas the Arr3-to-M_2_R ratio declined rapidly, approaching zero by 50 minutes (**Fig. 6B**). In constrast, the M_2_R concentration remained relatively stable, decreasing by ∼5% and ∼30% when co-expressed with Arr2 and Arr3, respectively, over the time course of stimulation. The difference in membrane dissociation dynamics between Arr2 and Arr3 is intriguing and warrants further investigation. One possible explanation is that activated Arr2 exhibits enhanced binding to PI(4,5)P_2_ in the membrane, whereas activated Arr3 shows reduced PI(4,5)P_2_ binding, which may facilitate its rapid membrane dissociation (**Fig. 2**).

FIF analysis on CCPs over the time course of carbachol stimulation revealed a gradual increase in both the Arr2-to-M_2_R and Arr3-to-M_2_R ratio within CCPs, indicating an elevated localization of Arr2 and Arr3 in CCPs as treatment time progresses (**Fig. 6C**). An overall higher Arr3-to-M_2_R ratio was observed over the 50-minute treatment period (**Fig. 6C**).

### Arr2 and Arr3 Migrate to CCPs Differently and Independently of M_2_R

M_2_R appears in the CCPs earlier than Arr2 and Arr3 (**Fig. 6A**), suggesting that M_2_R does not require Arr2 or Arr3 to enter CCPs and that M_2_R and arrestins likely migrate to the CCPs independently. To investigate this further, we measured the areas of CCPs containing M_2_R or Arr2/3, respectively, over the time course of carbachol stimulation (**Fig. 6D**). Overall, a larger CCP area contained M_2_R compared to Arr2 or Arr3. Arr3 appeared in the CCPs rapidly, and plateaued 10 minutes after ligand addition, whereas Arr2 increased in CCPs slowly (**Fig. 6D**).

We calculated the fraction of the CCP area containing both arrestins and M_2_R as a proxy for colocalization estimation (**Fig. 6E**). The results showed a similar trend as the area analysis, with CCPs containing both Arr3 and M_2_R increasing rapidly to plateau level within the first 10 minutes of stimulation, while CCPs containing both Arr2 and M_2_R showed a gradual, almost linear increase over the course of the treatment. Notably, even at the highest co-localization level, only ∼80% of CCPs contained both Arr2/3 and M_2_R, further suggesting that Arr2/3 are not essential for M_2_R entry into CCPs. Collectively, these results highlight the differences in CCP binding dynamics between these two highly homologous arrestin subtypes.

## Discussion

Arr2 and Arr3 share highly conserved sequences and structures; however, they differ in function^41–47^. Our findings suggest that differences in their membrane-binding properties may contribute to the functional specialization. The most surprising discovery is that Arr3 primarily relies on its finger loop, rather than the C-edge used by Arr2, for membrane interaction (**Fig. 1**). While the finger loop has been implicated in additional roles^20,24^, it is traditionally considered an activation sensor, inserting into the receptor core to ’sense’ its activation state—at least in Arr1^33,48^ and Arr2^16,21–23,49,50^. However, direct structural evidence supporting a similar role for Arr3 finger loop is lacking. In contrast, our recent structure of ACKR3 in complex with Arr3 reveals that the Arr3 finger loop inserts directly into the membrane^24^, consistent with our findings here. This raises the intriguing possibility that, rather than engaging the receptor core, the Arr3 finger loop may preferentially anchor to the surrounding membrane. This could lead to major differences in how Arr2 and Arr3 engage the same receptor, resulting in distinct functional outcomes. Further structural and functional studies are needed to test this hypothesis.

We showed that the membrane alone, regardless of lipid composition, does not promote the activation of Arr2 or Arr3 (**Fig. 3, 4**), as this would otherwise lead to non-specific activation of Arr2 and Arr3 in the absence of an activated and phosphorylated receptor. Notably, in the presence of an anchored phosphorylated peptide, the membrane greatly facilitates arrestin recruitment (**Extended Data Fig. 1B**) and potentiates its activation (**Fig. 4**). This effectively primes Arr2 and Arr3 to engage the nearby receptor core or the membrane, facilitating downstream signaling. It will be important to determine whether this effect is specific to ACKR3 due to its unique interaction with Arr2/3^24^, or if phosphorylated C-tails of other receptors with strong core interactions produce similar outcome.

Our study provides a comprehensive view of how the interplay between PI(4,5)P_2_ binding sites and other lipid-anchoring sites regulates the overall membrane binding of Arr2 and Arr3. Generally, PI(4,5)P_2_ serves as an additional anchor on the C-domain of both Arr2 and Arr3, enhancing their membrane interaction. However, for Arr3, when activated in absence of a phosphorylated receptor, PI(4,5)P_2_ has little or even adverse effects on membrane association (**Fig. 2**). This may result from a misalignment between the finger loop, the PI(4,5)P_2_ binding site, and the C-edge, which could impede membrane binding and facilitate dissociation or from global conformational changes to a state that is no longer favorable for lipid binding. This explains the rapid decline in Arr3 membrane presence observed in cells 30 minutes after M_2_R activation (**Fig. 6B**).

GPCR C-tails and intercellular loop 3 vary greatly in length, and different GRK subtypes have preferences for phosphorylation sites at specific distances from the receptor core^51–54^. Our study shows that the spacer length between the phosphorylation sites and membrane regulates the arrestin-membrane assembly (**Fig. 5**) and potentially the arrestin-receptor interaction. This suggests a previously unappreciated mechanism of regulation of arrestin signaling. Additionally, the combination of lipidated phosphopeptides and nanodiscs offers a powerful approach to investigate how the phosphorylation barcode influences arrestin functions.

FIF analysis allowed us to quantitatively track the localization of Arr2 and Arr3 at the plasma membrane and within the CCPs before and after M_2_R activation (**Fig. 6A-C**). Arr3 responds more rapidly to stimulation compared to Arr2, and Arr2 and Arr3 exhibit different membrane dissociation dynamics (**Fig. 6B**), which likely result from their contrasting reliance on PI(4,5)P₂ interaction after activation (**Fig. 2**). We also observed that M_2_R and arrestins entry the CCPs at different time points (**Fig. 6D, E**), consistent with the finding of Grimes et al^11^, which suggest that GPCRs and arrestins migrate to the CCPs independently.

The ability of two homologous non-visual arrestins to regulate the signaling of over 800 GPCRs underscores the importance of their distinct yet complementary membrane-binding mechanisms. Our study reveals a more significant role of membrane lipids in arrestin recruitment and activation than previously envisioned. We uncover unique membrane engagement strategies employed by Arr2 and Arr3, driven by differences in membrane-binding elements and contrasting reliance on PI(4,5)P_2_. These results highlight the critical role of the lipid environment in directing arrestin activation and demonstrate how receptor C-tail phosphorylation and membrane interactions collectively fine-tune arrestin function.

## Acknowledgements

We thank Dr. Vsevolod V. Gurevich in the Department of Pharmacology from Vanderbilt University for generous gifts of Arr2 and Arr3 plasmids and insightful comments on the manuscript. We thank Kathryn Schultz for her technical assistance with DEER data collection.

## Author contributions

Q.C. and V. R. conceptualize this study. Y. A, K. W. M. and Q.C. performed molecular cloning. K.W.M., Q.C., Y. A., S.Z., K.T., J.M.C. and R. T. expressed and purified arrestin variants. K.W.M., Q.C. and S.Z. performed the pulldown assays. Q.C., S.Z, and R.T. labelled arrestins and performed the fluorescence measurements. K. T. performed the trypsin digestion assay. Y.Z., C.K. and A.M. collected and analyzed the DEER data. T.D.K. and V.R. performed the cell FIF analysis. Q.C., Y.A., T.D.K. and V.R. wrote the original draft and all authors further edited the manuscript. Q.C., V.R., C.K., and A. M. contributed funding.

## Methods

### Expression and Purification of Arrestins

Expression and purification of Arr2/3 from *E. Coli* cells was described previously^55^. WT or variants of Arr2 and Arr3 were prepared using similar procedures. Briefly, the pTrcHisB plasmid containing bovine Arr2 or Arr3 was transformed into *E. coli* Rosetta cells and protein expression was induced with 25 µM (Arr2) or 37.5 µM (Arr3) IPTG for 4 hours at 30°C. The cell pellets were resuspended and homogenized in buffer containing 20 mM MOPS (pH 7.5), 400 mM NaCl, 5 mM EDTA, 2 mM DTT, 1 mM PMSF, leupeptin, and lima bean trypsin protease inhibitor. Cells were lysed using microfluidizer and the lysate was clarified by centrifugation at 18,000 rpm for 30 minutes. The supernatant was collected and arrestin was precipitated by the addition of (NH_4_)_2_SO_4_ to a final concentration 0.32 mg/ml. Precipitated arrestin was collected by centrifugation at 18,000 rpm for 60 minutes. The pellet was then dissolved in buffer containing 20 mM MOPS (pH 7.5), 2 mM EDTA, and 1 mM DTT, and then centrifuged at 18,000 rpm for 30 minutes to remove insoluble parts. The supernatant containing soluble arrestin was applied to a heparin column and eluted with a linear NaCl gradient (0.3-0.8 M for Arr2 and 0.5-1 M for Arr3). Fractions containing arrestin were identified by Coomassie blue staining and combined. For Arr2, the salt concentration of the pooled fractions was adjusted to 120 mM, loaded onto a 5 ml HiTrap Q HP column (Cytiva) and a 5 ml HiTrap SP column (Cytiva). Arr2 remains in the flowthrough while majority of the contamination bind to either Q or SP column. For Arr3, the salt concentration of the pooled fractions was adjusted to 100 mM, and the solution was loaded onto a linked 5 ml HiTrap Q HP (that it flows through) and 5 ml HiTrap SP HP column (that it binds). The columns were then uncoupled and a linear NaCl gradient (0.1-0.5 M) was used to elute Arr3 from the SP column. The fractions containing arrestin were concentrated with a 30 kDa cutoff Amico concentrator (Millipore) and further purified using a Superdex 200 increase 10/300 GL column (Cytiva) equilibrated with 20 mM MOPS (pH 7.5), 150 mM NaCl, and 0.5 mM TCEP. The peak fractions were collected, concentrated with a 30 kDa cutoff Amicon concentrator (Millipore), and stored at -80 °C.

Single Cysteine and double Cysteines were introduced to the Arr2 and Arr3 with all native cysteines mutated which retains its receptor-binding capacity^56^. Those variants were expressed and purified as described above with the following modifications. For Arr2-12C/392uring the size exclusion chromatograph purification step, 0.2 mM of DTT instead of 0.5 mM TCEP was used to avoid interfering with labeling. Fractions containing arrestin variants were concentrated with a 30 kDa concentrator.

### Expression and purification of NW9

Expression and purification of NW9 from *E. coli* cells was described previously^26^. Briefly, the pET28a plasmid containing NW9 was transformed into *E. coli* Rosetta cells under kanamycin selection, and protein expression was induced with 1mM IPTG at an OD^600^ of 0.6. Cells were incubated for another 3 hours at 37°C. A 4-liter cell pellet was resuspended in 50 mM HEPES (pH 8.0), 500 mM NaCl, 1% Triton-X 100, 1 mM PMSF, and 1mM leupeptin. Cells were lysed using a microfluidizer and the lysate was clarified by centrifugation at 12000 rpm for 30 minutes. The supernatant was collected and loaded twice at room temperature through 10mL HisPur^™^ Ni-NTA Resin (Thermo) that was equilibrated with 20 mM HEPES (pH 8.0) and 500 mM NaCl. The column was washed with 100 mL of 20 mM HEPES (pH 8.0), 500 mM NaCl, and 50 mM sodium cholate, followed by a second wash step with 100 ml of 20 mM HEPES (pH 8.0), 100 mM NaCl, and 20 mM imidazole. Elution of NW9 was carried out with 100 mL of 20 mM HEPES(pH 8.0), 100 mM NaCl, and 200 mM imidazole. Fractions containing NW9 were identified by coomassie blue staining of SDS-PAGE and combined. Combined fractions containing NW9 were concentrated with a 30 kDa cutoff Amico concentrator. ∼500 μL at a time of the concentrated fractions were loaded onto a 5 mL HiTrap Desalting column (GE) equilibrated with 50 mM HEPES (pH 8.0) and 100 mM NaCl. Fractions containing NW9 were concentrated to ∼10 mg/mL and stored at -80°C.

### Preparation of Nanodisc

The ratio between lipids and NW9 of 60 was used for assembling empty nanodiscs. POPC, POPS, POPG dissolved in chloroform and brain PI(4,5)P_2_ dissolved in chloroform, methanol and water were purchased from Avanti. POPS, POPG or PI(4,5)P_2_ were mixed with POPC at desired ratio and dried under N^2^ gas. The vials were left in a desiccator for at least 30 minutes to get rid of remaining chloroform. Sodium cholate was used to dissolve the lipids. NW9 and biobeads were added to the solubilized lipids and incubated on a rotator overnight at 4°C or three hours at room temperature.

### Nanodisc pulldown assays

Each binding assay reaction was set with a total volume of 50 μl containing 2 μM nanodisc and 2 μM Arr2 or Arr3 variants. The nanodisc and arrestin were incubated in the binding buffer containing 20 mM MOPS (pH7.5) and 100 mM NaCl for 20 minutes at room temperature. For each reaction, 3 μl of Ni-NTA magnetic beads (PureCube) was added and incubated on a rotator for 60 minutes at room temperature. For Arr3, 10 mM imidazole was included in the binding buffer to limit its nonspecific binding to Ni beads. The beads were washed with 1 ml of the binding buffer for three times and eluted with 20 μl of the binding buffer supplemented with 200 mM imidazole.

### Preparation of hyper-phosphorylated rhodopsin

The isolation of rod outer segment (ROS) and the preparation of hyper-phosphorylated rhodopsin was previously described^57^. In brief, the bovine retina was homogenized and resuspended with 25.5% sucrose. It was then overlaid on top of two sucrose layers containing 32.25% and 27.125% sucrose. The sucrose gradient was centrifuged in a swinging bucket rotor (SW-27) at 18,000 rpm for 90 minutes. ROS settled between the two layers and were resuspended in buffer containing 20 mM HEPES (pH 8.0) and 2 mM MgCl^2^. All procedures were conducted in a dark room under red light.

For phosphorylation, 2 μM of ROS was incubated 100 nM GRK1 in a buffer containing 50 mM HEPES (pH 8.0), 500 μM ATP, and 10 mM MgCl^2^ under light at room temperature for one hour. After phosphorylation, the membrane was washed twice with 20 mM HEPES (pH 8.0) and 100 mM NaCl and then incubated with 10-fold molar excess of 11-cis retinal in the dark at room temperature for one hour for regeneration.

### DEER Spectroscopy

The double Cys variant was incubated with a 40-fold molar excess of 4-maleimido-TEMPO (MAL-6, Sigma–Aldrich) overnight at 4 °C under gentle agitation. Excess spin labels were removed by dialysis with buffer containing 50 mM MOPS (pH 7.0) and 100 mM NaCl. DEER data were collected at 80K using a Q-band Bruker ELEXSYS 580 and an overcoupled EN5107D2 resonator as previously described^58^. Briefly, samples containing 20% deuterated glycerol as a cryoprotectant were held in quartz capillaries and flash-frozen in a dry ice and acetone mixture. Raw data were phased and background-corrected, plotted, and analyzed using the LongDistances software program written by C. Altenbach (University of California, Los Angeles, CA).

### Monobromobimane labeling of arrestin

Single Cys variants of Arr2 or Arr3 with all native cysteines mutated were expressed and purified as described above with the following modifications. During the Q/SP ion exchange purification step, reducing agents such as DTT was omitted in the elution buffer to avoid interfering with monobromobimane labeling. Fractions containing arrestin variants were concentrated to ∼5 mg/ml and twenty-molar excess of freshly prepared monobromobimane (Sigma) was added and incubated on ice overnight in the dark. The sample was loaded on a Superdex 200 Increase 10/300 GL column equilibrated with buffer containing 20 mM MOPS (pH 7.5), 150 mM NaCl and 0.5 mM TCEP to get rid of excess mBrB. The fractions containing mBrB-labelled arrestins were collected, concentrated with a 30 kDa cutoff Amicon concentrator, and stored at -80 °C. The mBrB labeling level was estimated to >75% based on the mBrB fluorescence absorption at 383 nm and the protein absorption at 280 nm.

### Bimane fluorescence measurements

The bimane-labelled Arr2 or Arr3 variant at 2 µM was incubated with 2 µM POPC, POPS, POPG or PI(4,5)P_2_ nanodisc at room temperature for 20 minutes. The reactions were transferred to a 384-well black plate (Corning) for fluorescence measurement using a plate reader (BioTek). The excitation wavelength was set to 375 nm and the absorption was monitored from 420 nm to 700 nm at step size of 2 nm. Each reaction was set in triplets and assays were repeated at least four times. For the assays with peptide/phosphopeptide, the reaction contained 0.5 µM bimane-labelled Arr2 or Arr3 variant, 0.5 µM nanodisc and 5 µM peptide/phosphopeptide.

### Limited trypsin digestion

Arr2 or Arr3 at 5 μM was incubated with 5 μM of p-*Rho, POPC, POPS, POPG, or PIP2 nanodiscs and 5 uM ACKR3_pp1_, ACKR3_pp2_, ACKR3_pp3_, ACKR3_pp4_, or buffer (20 mM MOPS, 100 mM NaCl, and 20 mM CaCl^2^) at room temperature for 10 minutes. Trypsin at a final concentration of 4 μg/ml was added to initiate the digestion at room temperature for 5 minutes. The reaction was then quenched with sample loading buffer.

### Cell culture and DNA transfection

HEK293 cells (ATCC) were grown in T25 flasks (CellTreat) in DMEM (high glucose) supplemented with 10% (v/v) fetal bovine serum, 100 U/ml penicillin, 100 mg/ml streptomycin. Cells were cultured in a humidified incubator with 95% air and 5% CO^2^ at 37°C. Two days prior to imaging, cells were plated in poly-d-lysine coated 35 mm dishes with a 14 mm #1.5 cover glass micro-well (Fisher Scientific) at a density of 2.0×10^5^ cells per dish. At 70-90% confluency the cells were transfected using Lipofectamine^TM^ 3000 reagent (Thermo Fisher Scientific) following manufacturers protocol. The DNA constructs were transfected at a 5:1 ratio (mEGFP-M_2_R to mCitrine-Arr2/3) to obtain optimal expression levels of both proteins. The cells were incubated in a humidified incubator for 24 hours post transfection. Prior to imaging, the transfected cells were removed from the incubator, washed with DPBS and suspended in DPBS.

### Configuration of two-photon micro-spectroscopy

This advanced setup allows for the separation of overlapping fluorescence signals from multiple fluorophores while reducing background fluorescence, thereby enabling accurate quantification of multiple fluorescently tagged proteins via pixel-level spectral unmixing. Fluoresce images (660×520 pixels^2^) were acquired using a two-photon optical micro-spectroscope incorporating a tunable femtosecond laser (MaiTai, Spectra Physics, Santa Clara, CA), a Zeiss Axio Observer inverted microscope stand, an OptiMiS detection head, a cooled electron-multiplying charge-coupled device (EMCCD) camera (iXonUltra 897, Andor Technologies), and a line-scan module^59^ from Aurora Spectral Technologies. The MaiTai laser was aligned to incorporate a spatial light modulator (SLM) (P1920-1152- HDMI Nematic SLM System, Meadowlark Optics) which was used to shape the laser into to a multi-beam array consisting of 24 individually focused beams as described previously ^60,61^. The multi-point array can excite 24 voxels in the sample simultaneously by delivering 100-fs pulses of 955 nm laser light through an infinity-corrected, C-Apochromat, water-immersion objective (x63, NA=1.2; Carl Zeiss Microscopy). The average power was 2.9 mW/voxel, the pixel dwell time was 5 ms, and the emission for each voxel was projected through a diffraction grating onto the EMCCD resulting in a spectral bandwidth of 450 to 600 nm with a spectral resolution of 11 nm.

### Cell imaging using two-photon micro-spectroscopy

A single field of view was imaged for each sample, selected to include at least two cells that express both M_2_R and arrestin. Five imaging scans were performed consecutively at different planes along the z-axis, in 0.5 µm increments, starting about 0.5 µm below the basolateral membrane. For each field of view, the image with the highest mEGFP-M_2_R signal, representing the plasma membrane, was selected for further analysis. Cells were imaged before and 2, 10, 20, 30, 40, and 50 minutes after 1 mM carbachol stimulation.

### Processing of fluorescence images

The fluorescence signals for each protein were quantified using an in-house software program, OptiMiS-DC. The emission spectra of EGFP and Citrine were determined from cells expressing only mEGFP-M_2_R or only mCitrine-Arr2/3 respectively. Those emission spectra were used to deconvolute (unmix) the composite spectrum collected from cells expressing both mEGFP-M_2_R and mCitrine-Arr2/3. Unmixing at the pixel level allows us to quantify the signal from individual protein^62^. The unmixed data were saved into separate two-dimensional fluorescence image stacks.

Once unmixed, the images were processed using an in-house software package, FIF spectrometry suite^63^. On the two-dimensional fluorescence image stack, regions of interest (ROIs) were hand-drawn within the boundaries of the basolateral membrane, excluding the outer edges, and further divided into segments containing ∼500 pixels each via a moving square segmentation method (**Extended Data Fig. 3A**)^60,64^. On average, each cell was divided to approximately 40 segments.

The fluorescence intensities of M_2_R and Arr2/3 at each pixel within a segment were used to generate two histograms whose means were used to calculate the concentrations of M_2_R and Arr2/3, respectively (**Extended Data Fig. 3A**). Fitting intensity histograms with a Gaussian curve at the segment level using the Nelder-Mead method^60,65,66^ effectively filters out extremely high or low intensity pixels, such as those originating from CCPs^67,68^, while accurately reflecting the average protein concentration on the uniform membrane. The mean (µ) and standard deviation (σ) of the Gaussian curve represent the mean intensity (〈𝐼_𝑠_〉) and the standard deviation (σ_s_) of the segment. Protein concentration is calculated at the segment level by inserting the mean intensity (〈𝐼_𝑠_〉) of each segment into Eq. 1, as expanded upon in the section below. This analysis was repeated for all segments in the unmixed images, effectively quantifying the individual concentration of M_2_R and arrestin at the segment level.

CCPs were analyzed separately from the uniform membrane by drawing ROIs around each pit individually on each unmixed image stack to quantify the formation of pits visible in the M_2_R maps and the arrestin maps respectively.

### Determination of protein concentrations

The M_2_R concentration at the plasma membrane were determined for individual segment within ROI by fitting the mean of the fitted Gaussian 〈𝐼_𝑠,𝑚𝑒𝑚_〉 to the following equation:

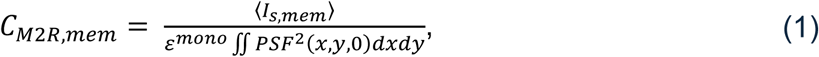

where 𝜀^𝑚𝑜𝑛𝑜^represents the brightness of a single fluorescent protein, PSF refers to the point spread function of the focused laser beam represented by a Gaussian-Lorentzian function^60^. For membrane proteins, such as M_2_R, the integral of the PSF is numerically evaluated at the basolateral membrane (𝑧 = 0).

The portions of arrestins located within the cytosol but near the membrane were also excited. To accurately determine the concentration of membrane-bound arrestin, we applied a correction to the concentration calculation, subtracting the contributions of cytosolic arrestin signal using the equation:

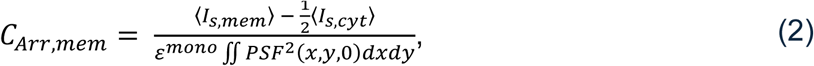

where, 〈𝐼_𝑠,𝑐𝑦𝑡_〉 represents the mean intensity of the segment in the cytosol; determined from image cross sections acquired 0.5-2 µm away from the basolateral membrane.

To determine 𝜀^𝑚𝑜𝑛𝑜^, fluorescent protein solutions (mEGFP and mCitrine)^69^ in HEPES buffer with 1 mM DTT were imaged on the same two-photon laser scanning micro-spectroscope used for cell imaging. Solutions of mEGFP and mCitrine were measured at multiple concentrations (6.2, 3.1, 1.6, 0.77, 0.39, 0.19 µM and 8.0, 4.0, 2.0, 1.0, 0.5, 0.25 µM respectively). The intensity data were plotted against respective molecular concentrations and fitted with a linear regression. The slope of the linear regression, Θ, was used to determine the monomeric brightness (𝜀^𝑚𝑜𝑛𝑜^) of each fluorescent protein using equation (S22) from ^60^, i.e.,

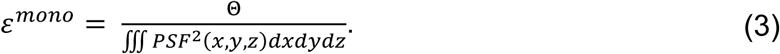

Concentrations were determined at the segment level (or ROI level in the case of CCPs) and the ratio between arrestin and M_2_R concentrations was calculated for every segment. The reason we used concentration ratios instead of separate concentrations for Arr2/3 and M_2_R was to cancel out systematic uncertainties inherent in the determination of concentrations in a volume (i.e., the excitation beam “volume”) that is poorly defined. Our protein concentration estimates are accurate up to a multiplicative constant, influenced by systematic errors related to sample area (or volume) and concentration variability across segments and cells. Since those constants are similar for M_2_R and arrestins, we analyzed the ratio of Arr2/3 to M_2_R concentrations in each segment. Data were separated by treatment time and a histogram of ratios was generated for each treatment time. Each histogram of ratios was then fitted with a Gaussian (normal) distribution and the mean (µ) and width (σ) were extracted for each treatment time (**Extended Data Fig. 3B, C**).

The data were bootstrapped using a resample with replacement method, where ROIs were sampled at random, allowing the same ROI to be selected multiple times. In a single bootstrapping event, the number of ROIs sampled was equal to the total number of ROIs in the dataset, and all data from the sampled ROIs were compiled into a histogram. The histogram was fitted to a Gaussian distribution which ranged from -5 to 5 with a bin size of 0.2. For display, plots were limited to a range of -4 to 4. After 100 repeated bootstrapping events the average and error of each fitting parameter was determined.

## Extended Data Figures

**Extended Data Fig. 1.**
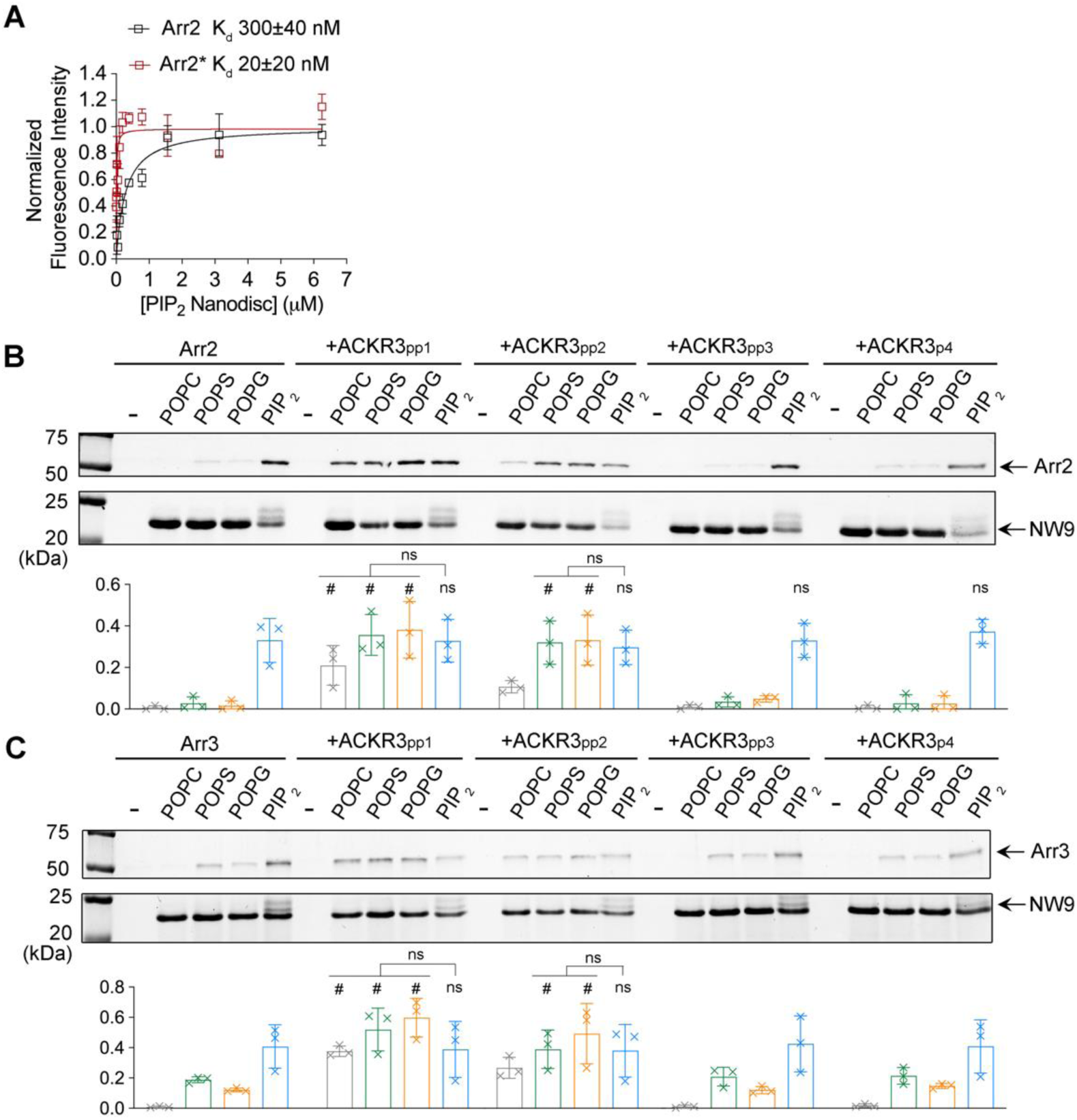
Preactivated Arr2 has increased binding affinity for the PI(4,5)P_2_ nanodiscs and Membrane tethered phosphopeptides anchor Arr2/3 at the membrane. **A.** Binding curves of Arr2 and Arr2* to the PI(4,5)P_2_ nanodiscs, determined by changes in the L338B bimane fluorescence intensity. **B, C.** A HIS pulldown assay shows that ACKR3_pp1_ and ACKR_pp2_, but not ACKR3_pp3_ and ACKR_p4_, promote Arr2 (**B**) and Arr3 (**C**) binding to the POPS, POPG nanodiscs at levels similar to those observed with the PI(4,5)P_2_ nanodiscs.

**Extended Data Fig. 2.**
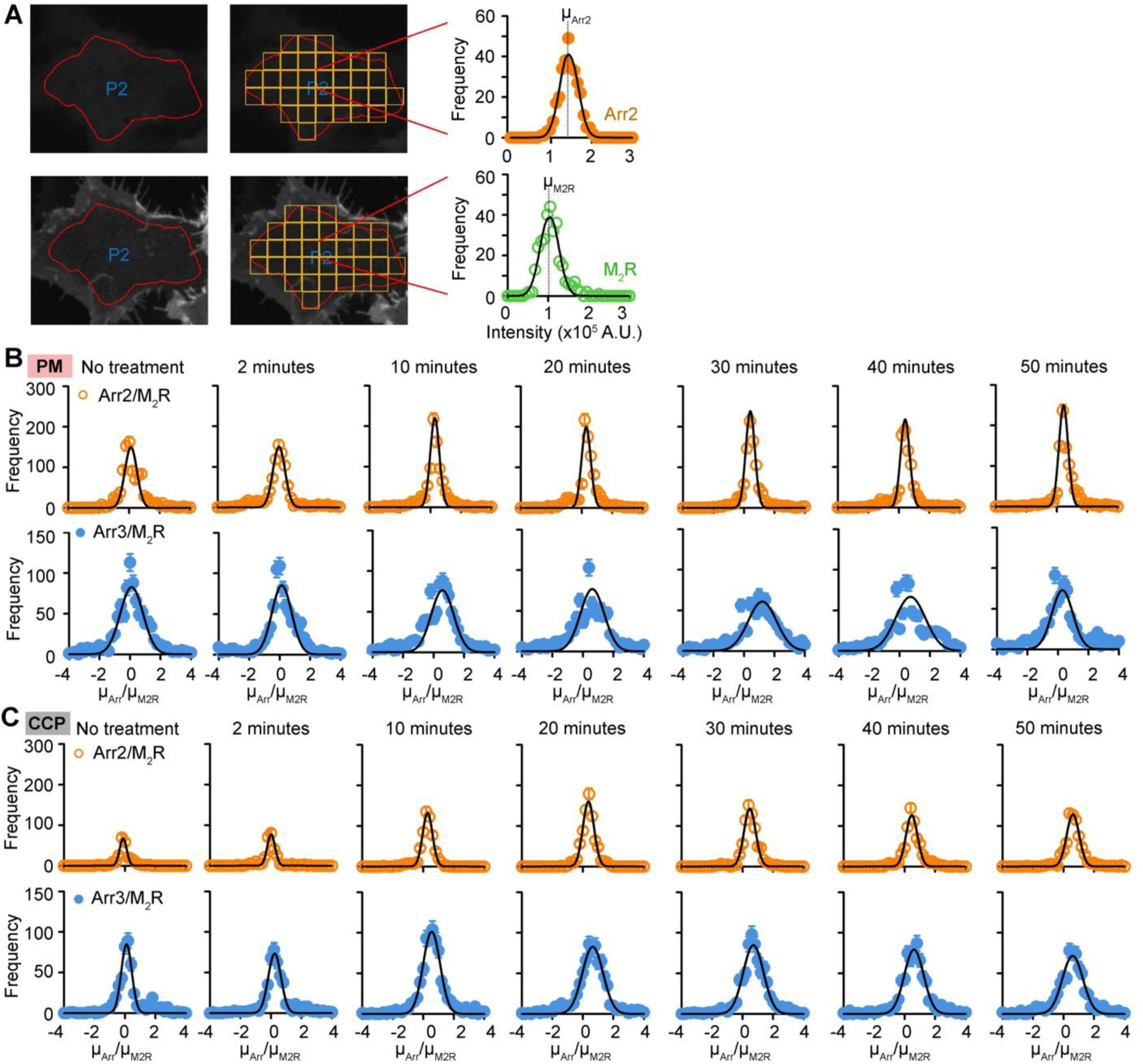
FIF analysis of Arr2/3 and M_2_R at the plasma membrane and within the CCPs before and after stimulation. **A.** ROIs were drawn using a polygon overlay tool (red outline), and each ROI was segmented into subsections (yellow boxes). Fluorescence intensities of M_2_R and Arr2 at each pixel were used to generate histograms for each segment, which were fitted with a Gaussian function. The mean and standard deviation of each the Gaussian curve was used to compute Arr2-to-M_2_R ratio for each segment in an image. This ratio computed from 300-500 segments were used to assemble histograms of Arr2-to-M_2_R ratios at each time point, which are shown in panel B. The process was repeated for samples co-expressing Arr3 and M_2_R. **B, C**. Distributions of Arr2/3-to-M_2_R concentration ratios at selected time points from uniform membrane (**B**) and CCP (**C**) analysis. The errors are the square root of the frequency of occurrence of each ROI segment. The histograms were fitted with a Gaussian to determine the mean of concentration ratios, *μ_Arr2/M2R_* and *μ_Arr3/M2R_*.

## References

1 Gurevich, V. V. & Gurevich, E. V. GPCR Signaling Regulation: The Role of GRKs and Arrestins. Front Pharmacol 10, 125 (2019). 10.3389/fphar.2019.00125

2 Luttrell, L. M. & Lefkowitz, R. J. The role of beta-arrestins in the termination and transduction of G-protein-coupled receptor signals. J Cell Sci 115, 455–465 (2002). 10.1242/jcs.115.3.455

3 Eiger, D. S. et al. Phosphorylation barcodes direct biased chemokine signaling at CXCR3. Cell Chem Biol 30, 362–382 e368 (2023). 10.1016/j.chembiol.2023.03.006

4 Kaya, A. I., Perry, N. A., Gurevich, V. V. & Iverson, T. M. Phosphorylation barcode-dependent signal bias of the dopamine D1 receptor. Proc Natl Acad Sci U S A 117, 14139–14149 (2020). 10.1073/pnas.1918736117

5 Yang, F. et al. Phospho-selective mechanisms of arrestin conformations and functions revealed by unnatural amino acid incorporation and (19)F-NMR. Nat Commun 6, 8202 (2015). 10.1038/ncomms9202

6 Bottke, T. et al. Exploring GPCR-arrestin interfaces with genetically encoded crosslinkers. EMBO Rep 21, e50437 (2020). 10.15252/embr.202050437

7 Aydin, Y. et al. Structural details of a Class B GPCR-arrestin complex revealed by genetically encoded crosslinkers in living cells. Nat Commun 14, 1151 (2023). 10.1038/s41467-023-36797-2

8 Gurevich, V. V. & Benovic, J. L. Visual arrestin interaction with rhodopsin. Sequential multisite binding ensures strict selectivity toward light-activated phosphorylated rhodopsin. J Biol Chem 268, 11628–11638 (1993).

9 Cahill, T. J., 3rd et al. Distinct conformations of GPCR-beta-arrestin complexes mediate desensitization, signaling, and endocytosis. Proc Natl Acad Sci U S A 114, 2562–2567 (2017). 10.1073/pnas.1701529114

10 Zhuang, T. et al. Involvement of distinct arrestin-1 elements in binding to different functional forms of rhodopsin. Proc Natl Acad Sci U S A 110, 942–947 (2013). 10.1073/pnas.1215176110

11 Grimes, J. et al. Plasma membrane preassociation drives beta-arrestin coupling to receptors and activation. Cell 186, 2238–2255 e2220 (2023). 10.1016/j.cell.2023.04.018

12 Lally, C. C., Bauer, B., Selent, J. & Sommer, M. E. C-edge loops of arrestin function as a membrane anchor. Nat Commun 8, 14258 (2017). 10.1038/ncomms14258

13 Janetzko, J. et al. Membrane phosphoinositides regulate GPCR-beta-arrestin complex assembly and dynamics. Cell 185, 4560–4573 e4519 (2022). 10.1016/j.cell.2022.10.018

14 Chen, Q. & Tesmer, J. J. G. G protein-coupled receptor interactions with arrestins and GPCR kinases: The unresolved issue of signal bias. J Biol Chem 298, 102279 (2022). 10.1016/j.jbc.2022.102279

15 Zhai, R. et al. Distinct activation mechanisms of beta-arrestin-1 revealed by (19)F NMR spectroscopy. Nat Commun 14, 7865 (2023). 10.1038/s41467-023-43694-1

16 Huang, W. et al. Structure of the neurotensin receptor 1 in complex with beta-arrestin 1. Nature 579, 303–308 (2020). 10.1038/s41586-020-1953-1

17 Kim, K. & Chung, K. Y. Molecular mechanism of beta-arrestin-2 pre-activation by phosphatidylinositol 4,5-bisphosphate. EMBO Rep 25, 4190–4205 (2024). 10.1038/s44319-024-00239-x

18 Chen, Q. et al. Structural basis of arrestin-3 activation and signaling. Nat Commun 8, 1427 (2017). 10.1038/s41467-017-01218-8

19 Milano, S. K., Kim, Y. M., Stefano, F. P., Benovic, J. L. & Brenner, C. Nonvisual arrestin oligomerization and cellular localization are regulated by inositol hexakisphosphate binding. J Biol Chem 281, 9812–9823 (2006). 10.1074/jbc.M512703200

20 Chen, K. et al. Tail engagement of arrestin at the glucagon receptor. Nature 620, 904–910 (2023). 10.1038/s41586-023-06420-x

21 Bous, J. et al. Structure of the vasopressin hormone-V2 receptor-beta-arrestin1 ternary complex. Sci Adv 8, eabo7761 (2022). 10.1126/sciadv.abo7761

22 Staus, D. P. et al. Structure of the M2 muscarinic receptor-beta-arrestin complex in a lipid nanodisc. Nature 579, 297–302 (2020). 10.1038/s41586-020-1954-0

23 Lee, Y. et al. Molecular basis of beta-arrestin coupling to formoterol-bound beta1-adrenoceptor. Nature 583, 862–866 (2020). 10.1038/s41586-020-2419-1

24 Chen, Q. et al. ACKR3–arrestin2/3 complexes reveal molecular consequences of GRK-dependent barcoding. bioRxiv, 2023.2007.2018.549504 (2023). 10.1101/2023.07.18.549504

25 Baidya, M. et al. Key phosphorylation sites in GPCRs orchestrate the contribution of beta-Arrestin 1 in ERK1/2 activation. EMBO Rep 21, e49886 (2020). 10.15252/embr.201949886

26 Nasr, M. L. et al. Covalently circularized nanodiscs for studying membrane proteins and viral entry. Nat Methods 14, 49–52 (2017). 10.1038/nmeth.4079

27 Gaidarov, I., Krupnick, J. G., Falck, J. R., Benovic, J. L. & Keen, J. H. Arrestin function in G protein-coupled receptor endocytosis requires phosphoinositide binding. EMBO J 18, 871–881 (1999). 10.1093/emboj/18.4.871

28 Sommer, M. E., Smith, W. C. & Farrens, D. L. Dynamics of arrestin-rhodopsin interactions: arrestin and retinal release are directly linked events. J Biol Chem 280, 6861–6871 (2005). 10.1074/jbc.M411341200

29 Vishnivetskiy, S. A. et al. An additional phosphate-binding element in arrestin molecule. Implications for the mechanism of arrestin activation. J Biol Chem 275, 41049–41057 (2000). 10.1074/jbc.M007159200

30 He, K. et al. Dynamics of phosphoinositide conversion in clathrin-mediated endocytic traffic. Nature 552, 410–414 (2017). 10.1038/nature25146

31 Kim, K. & Chung, K. Y. Molecular mechanism of beta-arrestin-2 pre-activation by phosphatidylinositol 4,5-bisphosphate. EMBO Rep (2024). 10.1038/s44319-024-00239-x

32 Schafer, C. T., Chen, Q., Tesmer, J. J. G. & Handel, T. M. Atypical Chemokine Receptor 3 "Senses" CXC Chemokine Receptor 4 Activation Through GPCR Kinase Phosphorylation. Mol Pharmacol 104, 174–186 (2023). 10.1124/molpharm.123.000710

33 Kang, Y. et al. Crystal structure of rhodopsin bound to arrestin by femtosecond X-ray laser. Nature 523, 561–567 (2015). 10.1038/nature14656

34 Kumari, P. et al. Functional competence of a partially engaged GPCR-beta-arrestin complex. Nat Commun 7, 13416 (2016). 10.1038/ncomms13416

35 Thomsen, A. R. B. et al. GPCR-G Protein-beta-Arrestin Super-Complex Mediates Sustained G Protein Signaling. Cell 166, 907–919 (2016). 10.1016/j.cell.2016.07.004

36 Kumari, P. et al. Core engagement with beta-arrestin is dispensable for agonist-induced vasopressin receptor endocytosis and ERK activation. Mol Biol Cell 28, 1003–1010 (2017). 10.1091/mbc.E16-12-0818

37 Lambert, L. et al. Endocytosis of Activated Muscarinic m2 Receptor (m2R) in Live Mouse Hippocampal Neurons Occurs via a Clathrin-Dependent Pathway. Front Cell Neurosci 12, 450 (2018). 10.3389/fncel.2018.00450

38 Roth, M. G. Phosphoinositides in constitutive membrane traffic. Physiol Rev 84, 699–730 (2004). 10.1152/physrev.00033.2003

39 Stoneman, M. R. et al. A general method to quantify ligand-driven oligomerization from fluorescence-based images. Nat Methods 16, 493–496 (2019). 10.1038/s41592-019-0408-9

40 Killeen, T. D. et al. Fluorescence Intensity Fluctuation Analysis of Protein Oligomerization in Cell Membranes. Curr Protoc 2, e384 (2022). 10.1002/cpz1.384

41 Morris, G. E. et al. Arrestins 2 and 3 differentially regulate ETA and P2Y2 receptor-mediated cell signaling and migration in arterial smooth muscle. Am J Physiol Cell Physiol 302, C723–734 (2012). 10.1152/ajpcell.00202.2011

42 Ahn, S., Wei, H., Garrison, T. R. & Lefkowitz, R. J. Reciprocal regulation of angiotensin receptor-activated extracellular signal-regulated kinases by beta-arrestins 1 and 2. J Biol Chem 279, 7807–7811 (2004). 10.1074/jbc.C300443200

43 Carey, L. M., Rice, R. J. & Prus, A. J. The Neurotensin NTS(1) Receptor Agonist PD149163 Produces Antidepressant-Like Effects in the Forced Swim Test: Further Support for Neurotensin as a Novel Pharmacologic Strategy for Antidepressant Drugs. Drug Dev Res 78, 196–202 (2017). 10.1002/ddr.21393

44 Oneda, B. et al. beta-Arrestin2 influences the response to methadone in opioid-dependent patients. Pharmacogenomics J 11, 258–266 (2011). 10.1038/tpj.2010.37

45 Sun, D., Ma, J. Z., Payne, T. J. & Li, M. D. Beta-arrestins 1 and 2 are associated with nicotine dependence in European American smokers. Mol Psychiatry 13, 398–406 (2008). 10.1038/sj.mp.4002036

46 Ikeda, M. et al. Possible association of beta-arrestin 2 gene with methamphetamine use disorder, but not schizophrenia. Genes Brain Behav 6, 107–112 (2007). 10.1111/j.1601-183X.2006.00237.x

47 Bohn, L. M. et al. Enhanced morphine analgesia in mice lacking beta-arrestin 2. Science 286, 2495–2498 (1999). 10.1126/science.286.5449.2495

48 Vishnivetskiy, S. A., Huh, E. K., Gurevich, E. V. & Gurevich, V. V. The finger loop as an activation sensor in arrestin. J Neurochem 157, 1138–1152 (2021). 10.1111/jnc.15232

49 Cao, C. et al. Signaling snapshots of a serotonin receptor activated by the prototypical psychedelic LSD. Neuron (2022). 10.1016/j.neuron.2022.08.006

50 Yin, W. et al. A complex structure of arrestin-2 bound to a G protein-coupled receptor. Cell Res 29, 971–983 (2019). 10.1038/s41422-019-0256-2

51 Nobles, K. N. et al. Distinct phosphorylation sites on the beta(2)-adrenergic receptor establish a barcode that encodes differential functions of beta-arrestin. Sci Signal 4, ra51 (2011). 10.1126/scisignal.2001707

52 Kawakami, K. et al. Heterotrimeric Gq proteins act as a switch for GRK5/6 selectivity underlying beta-arrestin transducer bias. Nat Commun 13, 487 (2022). 10.1038/s41467-022-28056-7

53 Drube, J. et al. GPCR kinase knockout cells reveal the impact of individual GRKs on arrestin binding and GPCR regulation. Nat Commun 13, 540 (2022). 10.1038/s41467-022-28152-8

54 Underwood, O. et al. Key phosphorylation sites for robust beta-arrestin2 binding at the MOR revisited. Commun Biol 7, 933 (2024). 10.1038/s42003-024-06571-1

## Method References

55 Vishnivetskiy, S. A., Zhan, X., Chen, Q., Iverson, T. M. & Gurevich, V. V. Arrestin expression in E. coli and purification. Curr Protoc Pharmacol 67, 2 11 11–12 11 19 (2014). 10.1002/0471141755.ph0211s67

56 Zhuo, Y., Vishnivetskiy, S. A., Zhan, X., Gurevich, V. V. & Klug, C. S. Identification of receptor binding-induced conformational changes in non-visual arrestins. J Biol Chem 289, 20991–21002 (2014). 10.1074/jbc.M114.560680

57 Chen, Q. et al. Structures of rhodopsin in complex with G-protein-coupled receptor kinase 1. Nature 595, 600–605 (2021). 10.1038/s41586-021-03721-x

58 Zhuo, Y., Gurevich, V. V., Vishnivetskiy, S. A., Klug, C. S. & Marchese, A. A non-GPCR-binding partner interacts with a novel surface on beta-arrestin1 to mediate GPCR signaling. J Biol Chem 295, 14111–14124 (2020). 10.1074/jbc.RA120.015074

59 Biener, G. et al. Development and Experimental Testing of an Optical Micro-Spectroscopic Technique Incorporating True Line-Scan Excitation. International Journal of Molecular Sciences 15, 261–276 (2014). 10.3390/ijms15010261

60 Stoneman, M. R. et al. A general method to quantify ligand-driven oligomerization from fluorescence-based images. Nature Methods 16, 493–496 (2019). 10.1038/s41592-019-0408-9

61 Stoneman, M. R. Y., K; Biener, G.; Killeen, T. D.; Rahman, S.; Harikumar, K.; Miller, L. J.; Raicu, V. Mechanistic insights from the atomic-level quaternary structure of short-lived GPCR oligomers in live cells. Under review (Nature Portfolio) (2024). 10.21203/rs.3.rs-4683780/v1

62 Patowary, S. et al. Experimental verification of the kinetic theory of FRET using optical microspectroscopy and obligate oligomers. Biophys J 108, 1613–1622 (2015). 10.1016/j.bpj.2015.02.021

63 Killeen, T. D. et al. Fluorescence Intensity Fluctuation Analysis of Protein Oligomerization in Cell Membranes. Current Protocols 2, e384 (2022). 10.1002/cpz1.384

64 Stoneman, M. R., Raicu, N., Biener, G. & Raicu, V. Fluorescence-based Methods for the Study of Protein-Protein Interactions Modulated by Ligand Binding. Curr Pharm Des 26, 5668–5683 (2020). 10.2174/1381612826666201116120934

65 Nelder, J. A. & Mead, R. A simplex method for function minimization. Computer Journal 7, 308–313 (1965). 10.1093/comjnl/7.4.308

66 Lagarias, J. C., Reeds, J. A., Wright, M. H. & Wright, P. E. Convergence properties of the Nelder-Mead simplex method in low dimensions. Siam Journal on Optimization 9, 112–147 (1998). 10.1137/S1052623496303470

67 Biener, G., Stoneman, M. R. & Raicu, V. Fluorescence intensity fluctuation analysis of receptor oligomerization in membrane domains. Biophys J 120, 3028–3039 (2021). 10.1016/j.bpj.2021.06.015

68 Stoneman, M. R., Biener, G. & Raicu, V. Reply to: Spatial heterogeneity in molecular brightness. Nat Methods 17, 276–278 (2020). 10.1038/s41592-020-0735-x

69 Adhikari, D. P. et al. Comparative photophysical properties of some widely used fluorescent proteins under two-photon excitation conditions. Spectrochimica Acta Part A: Molecular and Biomolecular Spectroscopy 262, 120133 (2021). 10.1016/j.saa.2021.120133

